# Olfactory-mediated host tracking and rapid local adaptation in a seed-feeding wasp

**DOI:** 10.64898/2026.10.01.755887

**Authors:** Xiaoxiao Chen, Y. Miles Zhang, Jingge Kuang, Yi Wang, Jialiang Li, Wenqian Hu, Senlin Yang, Wentao Wang, Hongliang Lv, Tao Li, Graham N. Stone, Kangshan Mao

**Author notes:** These authors contributed equally to this work (Xiaoxiao Chen, Y. Miles Zhang, Jingge Kuang). Correspondence (Kangshan Mao), (Graham N. Stone).

## Abstract

Understanding how specialized herbivores track divergence among their hosts is central to evolutionary biology. Congruent genetic structure between hosts and herbivores is often interpreted as evidence of long-term codivergence, but similar patterns may also arise when herbivores rapidly adapt to pre-existing host differentiation. Here, we integrate population genomics, host volatile profiling, and behavioral assays to examine host-associated divergence in the obligate seed-feeding wasp *Megastigmus daduheensi*s and its conifer host *Cupressus chengiana* across three river valleys on the eastern Qinghai-Tibet Plateau. Despite strong phylogeographic congruence between each of the three wasp and host lineages, calibrated demographic analyses indicate that wasp divergence occurred during the early Holocene, substantially later than the Pliocene-Pleistocene divergence of host lineages. This temporal mismatch is inconsistent with strict long-term codivergence and instead supports rapid host tracking in a topographic complex region. Host lineages differed in volatile organic compound profiles, and behavioral assays showed that wasps were preferentially attracted to volatiles from their local host lineages. These results identify olfactory-mediated host recognition as a plausible mechanism promoting host fidelity and reinforcing host-associated genomic and behavioral differentiation. Our findings show that apparently concordant host-herbivore divergence can emerge over compressed evolutionary timescales through rapid local adaptation, highlighting how ecological differentiation can accumulate within species through host tracking.

**Significance Statement:** Congruent genetic structure between specialized herbivores and their hosts is often taken as evidence of long-term codivergence. Combining population genomics, host volatile profiling, and behavioral assays, we show that closely matched host-herbivore divergence can instead arise when a seed-feeding wasp rapidly tracks pre-existing divergence among conifer host lineages. Estimated wasp divergence occurred in the Holocene-thousands rather than millions of years ago-substantially later than host divergence, and wasps preferentially responded to volatile profiles from their local hosts. These findings identify olfactory-mediated host recognition as a mechanism promoting host fidelity and rapid local adaptation, and caution against inferring long-term codivergence from matching genetic structure alone.

## Main

Understanding how local adaptation and population divergence arise remains a long-standing problem in evolutionary biology, because these processes shape the accumulation of ecological differentiation within species and may set the stage for deeper evolutionary divergence^1, 2^. Plant-feeding insects are particularly informative in this context. Their population structure can be shaped by host identity, host chemistry, host-associated habitats, and local environmental heterogeneity^3, 4^. When specialized herbivores and their host plants show concordant genetic or phylogeographic structure, this pattern is often interpreted as evidence of long-term codivergence^4-7^. However, such congruence is not necessarily diagnostic of synchronous evolutionary histories. It can also emerge when herbivores track pre-existing divergence among host lineages after host differentiation has already occurred^8-10^. In this case, herbivore divergence may be much younger than host divergence, even though the resulting population structure appears closely matched between trophic levels. Distinguishing long-term codivergence from recent host tracking therefore requires explicit comparison of divergence times, together with evidence for the ecological mechanisms that maintain host-associated differentiation.

Host tracking in specialist herbivores is likely to depend, at least in part, on sensory cues used during host recognition and host choice. Volatile organic compounds (VOCs) are among the most important cues mediating plant-insect interactions, and variation in volatile blends can influence feeding, mating, and oviposition decisions^11, 12^. In conifers, chemical traits also play a central role in defense and in interactions with specialized insect associates, including seed and cone-feeding taxa^13, 14^. Although host-associated divergence has been examined in several well-studied plant-insect systems, including figs, gall-inducing insects, and oak gall wasps^6, 8, 15, 16^, conifer seed-feeders remain less frequently used to test how host chemical differentiation contributes to fine-scale herbivore divergence.

Here we examine this question in the obligate seed-feeding wasp *Megastigmus daduheensis* and its conifer host *Cupressus chengiana* across three river valleys on the eastern Qinghai-Tibet Plateau^17, 18^. The host species *C. chengiana* has undergone allopatric divergence of two lineages and hybrid origin of the third lineage in the northern Hengduan Mountains (HDMs), a region whose biogeographic history has been shaped by mountain uplift and Quaternary climatic change^19-21^. This system therefore allows us to ask whether genetic congruence between a specialist seed-feeder and its host reflects synchronous codivergence or more recent tracking of host population divergence. We combined population genomic analyses, demographic modelling, host volatile profiling, and behavioral assays to address three questions: (1) Do wasp populations show genetic structure that corresponds to host lineages across river valleys? (2) Does the timing of wasp divergence support synchronous codivergence or later host tracking? (3) Do differences in seed-cone VOCs and wasp behavioral responses provide a mechanism by which host fidelity and host-associated differentiation could be maintained?

We found that *M. daduheensis* populations are genetically structured in a pattern that closely parallels host lineages, but their divergence is estimated to have occurred during the Holocene, substantially later than host divergence. This temporal asymmetry is inconsistent with strict long-term codivergence and is more consistent with rapid host tracking at a microgeographic scale. Host lineages differed in seed-cone VOC profiles, and wasps showed preferential behavioral responses to volatiles from their local host lineages. These results suggest that olfactory-mediated host recognition may promote host fidelity and contribute to host-associated genomic and behavioral differentiation. More broadly, our study shows that close correspondence between host and herbivore population structure can arise over short evolutionary timescales through local adaptation, without requiring synchronous codivergence between trophic levels.

## Results

### Sampling and genomic variation in the wasp system

We sampled 211 wasps from 11 populations over the three target river catchments Daduhe (DDH), Minjiang (MJR) and Bailong (BLJ) in the HDMs (**Fig. 1 and Supplementary Table 1**). The three river catchments are separated by approximately 50–150 km, a spatial scale over which dispersal may be limited in *M. daduheensis*, given its small body size (2–4 mm) and likely reliance on passive or short-distance movement. Whole-genome resequencing yielded 297 Gb of data, representing an average genome alignment depth of 18.4× (**Supplementary Table 2**). We identified 691,670 high-quality SNPs after quality control.

**Fig. 1.**
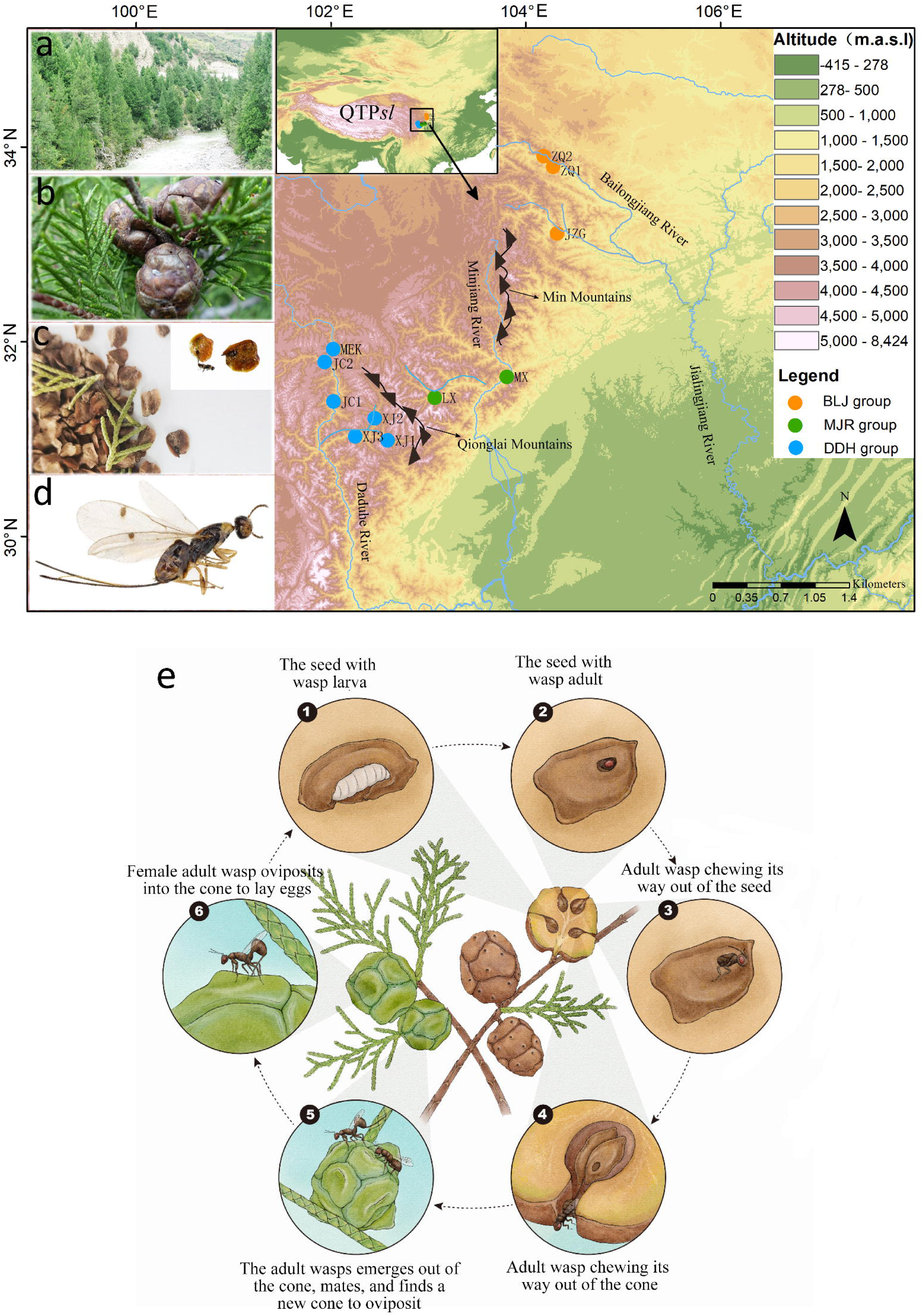
| Geographical distributions of sampled *M. daduheensis* and *C. chengiana* populations. QTPsl is the abbreviation of Qinghai-Tibet Plateau *sensu lato*. **a**, The habitat of *C. chengiana.* b, Cones of *C. chengiana* infected with wasps. c, *M. daduheensis* that lives inside the seeds. **d,** The lateral habitus of an adult female *M. daduheensis.* e, The life cycle of *M. daduheensis* in the cones and seeds of *C. chengiana*.

For analyses sensitive to linkage among markers, including ADMIXTURE and PCA, we used LD-pruned SNP datasets comprising 188,938 SNPs in *M. daduheensis* and 136,286 SNPs in *C. chengiana*. Demographic analyses were based on separately constructed datasets described in Methods, with callable genomic sites and invariant sites treated explicitly where required.

### Wasp and host lineages show concordant river-valley structure

ADMIXTURE analyses indicated that three genetic groups (*K* = 3) were optimal for both the wasps and the hosts (**Fig. 2a, b and Supplementary Fig. 1**). These three genetic groups corresponded closely to the DDH, MJR, and BLJ river catchments in both species. At lower *K* values, population structure was less fully resolved, but the optimal *K* = 3 result, PCA, and phylogenetic reconstruction all supported three river-valley-associated lineages in both trophic levels. PCA separated individuals from the three catchments in both *M. daduheensis* and *C. chengiana* (**Fig. 2c, d**), and maximum-likelihood phylogenetic analyses recovered reciprocally monophyletic valley lineages in both species (**Fig. 2e, f**). In both trees, MJR and BLJ were inferred as sister lineages relative to DDH.

**Fig. 2.**
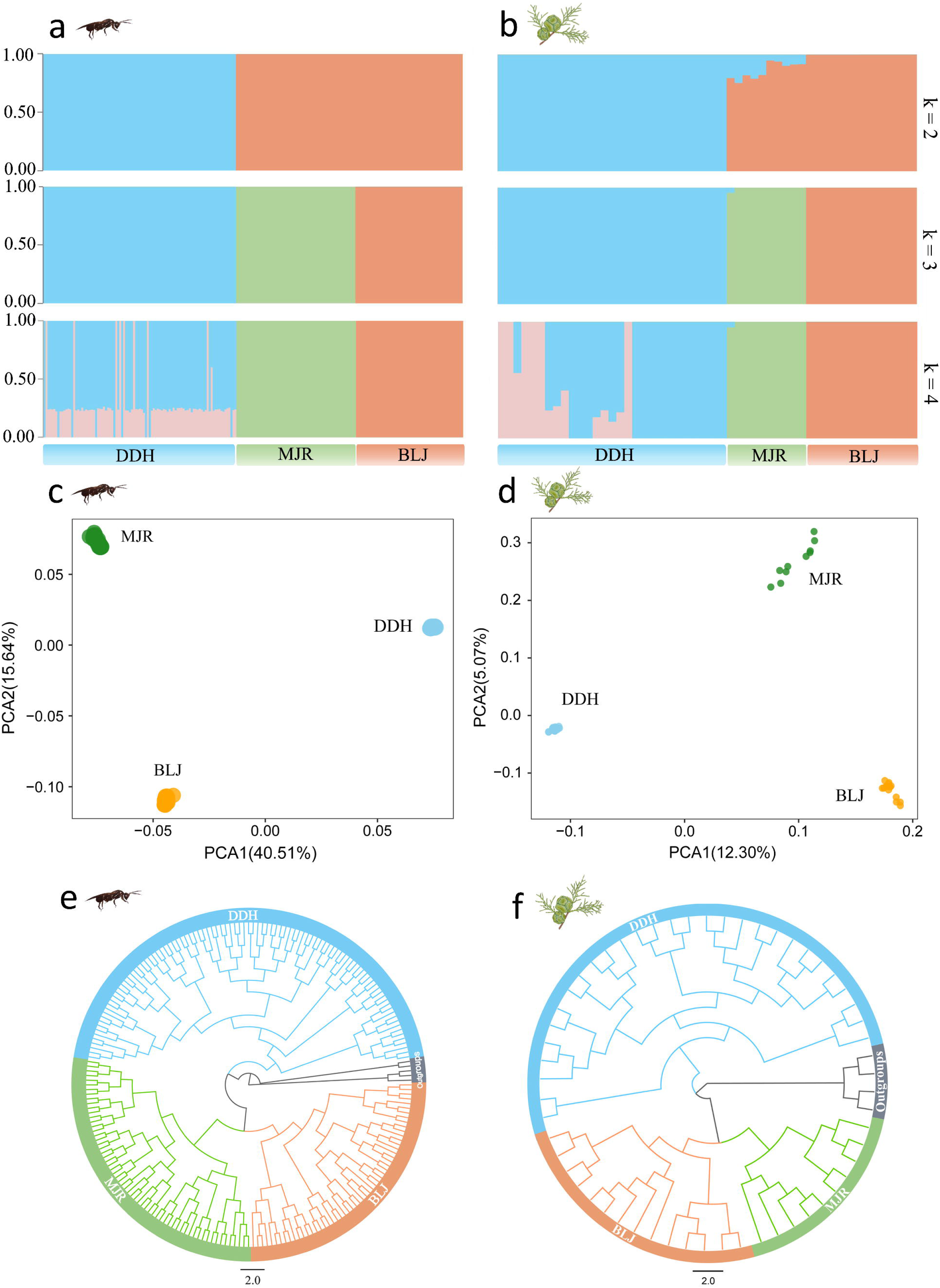
| Co-structure of *M. daduheensis* and *C. chengiana*. **a, b,** Admixture proportions of genetic clusters of different groups for *K* values of 2, 3 and 4 (with an optimal *K* = 3). **c, d,** Principal component analysis (PCA) plot for the 211 wasp and 53 host individuals based on the first two principal components. **e, f,** Maximum likelihood (ML) trees based on 211 wasp and 53 cypress individuals with outgroups using IQ-TREE. The node support rates of the three main branches are all 100. In all diagram parts, blue=DDH, green = MJR and red=BLJ.

Measures of genetic differentiation and diversity had similar trends between the wasps and the hosts (**Supplementary Fig. 2a–d and Supplementary Table 3**), which are represented by the subscript abbreviations of *M.D.* (for *M. daduheensis*) and *C.C.* (for *C. chengiana*), respectively. Population genetic distance was highest between BLJ and DDH (*F_st-M.D_* = 0.4930, *F_st-C.C_* = 0.2442). The average *π* value of DDH was lowest (*π_-M.D_* = 0.00027, *π_-C.C_* = 0.00191), while BLJ (*π_-M.D_* = 0.00039, *π_-C.C_* = 0.00229) was less than that of MJR (*π_-M.D_* = 0.00045, *π_-C.C_* = 0.00256). Other summary statistics, including Tajima’s *D* and LD decay, are reported in Supplementary Fig. 2 and Supplementary Table 3. DDH exhibits the slowest LD decay, MJR the fastest, while BLJ falls in between. Because LD decay can reflect multiple demographic and genomic processes, we do not use it alone as evidence for reduced genetic diversity.

### Wasp divergence substantially postdates host divergence

We inferred the demographic history of the three *M. daduheensis* lineages using fastsimcoal2 and SMC++. Among the eight fastsimcoal2 models tested, model 2 and model 8 received the strongest support (**Supplementary Table 4 and Supplementary Fig. 4**). Model 2 assumes a sequential divergence topology in which MJR and BLJ are sister lineages relative to DDH, whereas model 8 represents a simultaneous radiation of the three populations. Although model 8 had the lowest AIC, SMC++ independently recovered a sequential rather than simultaneous radiation pattern, consistent with model 2. Because both models inferred very recent wasp divergence relative to host divergence, the main conclusion of temporal asymmetry was robust to this topological uncertainty. We therefore present model 2 as the primary demographic scenario, while retaining model 8 as an alternative model in the Supplementary Information.

Under model 2, the split separating DDH from the MJR/BLJ lineage was estimated at approximately 9.10 thousand years ago (Kya; 95% CI, 4.21-167.54 Kya), and the MJR-BLJ split was estimated at 7.8 Kya (95% CI, 3.6-134.7 Kya) (**Fig. 3a and Supplementary Table 5**). Thus, the point estimates place wasp divergence in the Holocene, although the confidence intervals extend into the late Pleistocene. Importantly, even the upper bounds of these confidence intervals remain far younger than the earliest divergence inferred for the host. Significant asymmetric gene flow was inferred among the DDH, MJR, and BLJ wasp lineages. Gene flow was higher between BLJ and MJR than between either of these lineages and DDH.

**Fig. 3.**
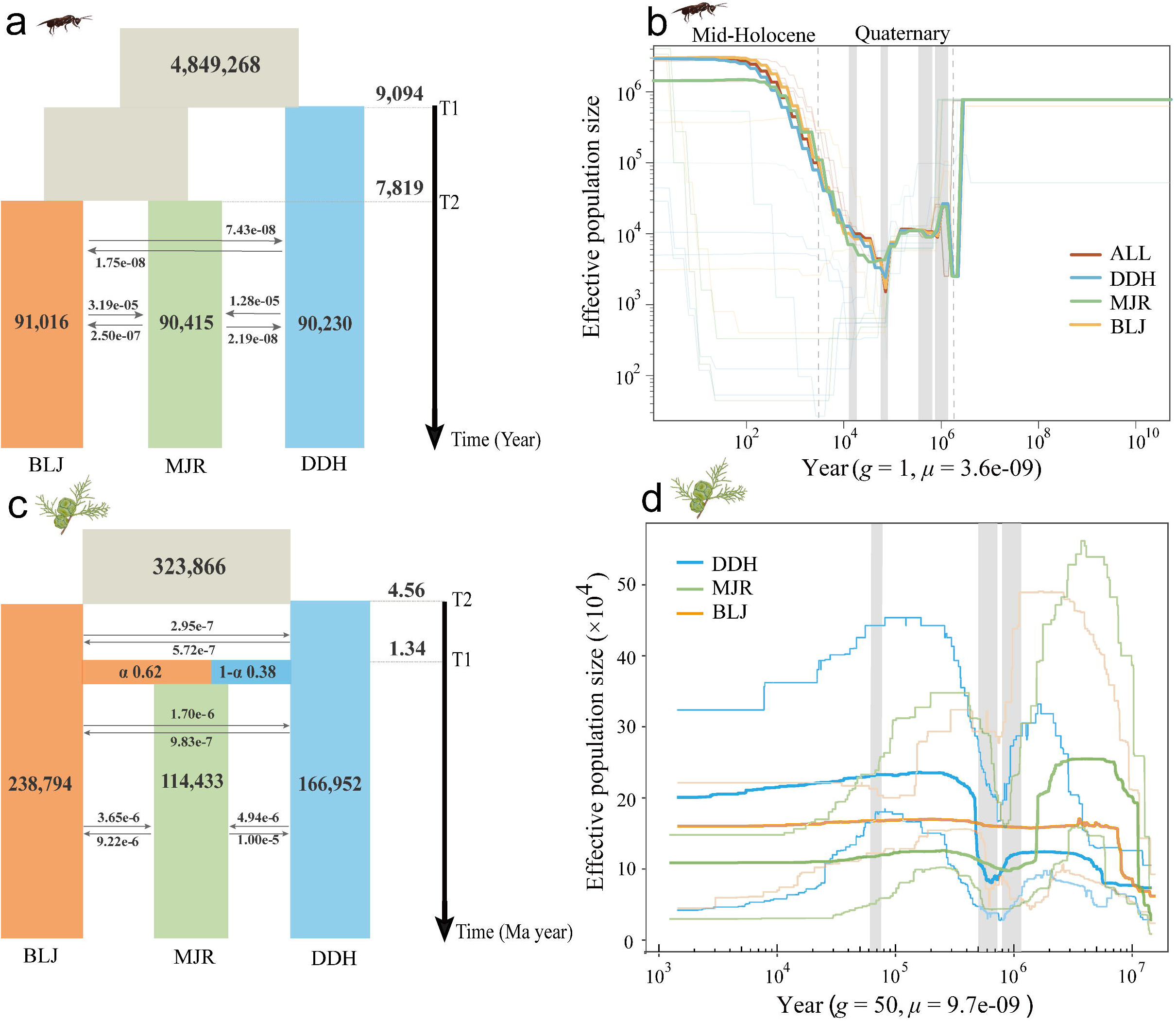
| Demographic history of *M. daduheensis* and *C. chengiana*. **a, c,** Maximum likelihood parameter estimates for the best fit (secondary contact model) demographic scenarios for *M. daduheensis* (**a**) and *C. chengiana* (**c**) in *fastsimcoal2*. Numbers within population bars indicate effective population sizes while numbers above/below arrows indicate the per generation migration rate between lineages. Numbers by the temporal scale indicate ages for population divergence or hybridisation. **b,** Historical effective population size in the different lineages and predicted by SMC++. The gray dotted lines depict the onset of the Quaternary and of the middle Holocene. The periods of the Xixiabangma Glaciation, the Naynayxungla Glaciation, the Marine Isotope Stage 5b (MIS5b) and the last glacial maximum (LGM) are highlighted with gray vertical bars from right to left. **d,** The detailed population demographic history of *C. chengiana* over the last 10 million years inferred by the Stairway Plot method ^24^. The thick lines represent the median, and the thin light lines represent the 95% pseudo-CI defined by the 2.5% and 97.5% estimations from the SFS analysis. The periods of the Xixiabangma Glaciation, the Naynayxungla Glaciation and MIS5b are highlighted with gray vertical bars from right to left^43^.

In contrast, divergence among *C. chengiana* lineages was inferred to be orders of magnitude older than divergence among wasp lineages (**Fig. 3c, d and Supplementary Fig. 3d**). The BLJ and DDH host lineages first diverged around 4.56 Mya (95% CI, 4.23–4.87 Mya), and the MJR lineage was inferred to have originated through hybridization around 1.34 Mya (95% CI, 1.14–1.41 Mya). Even when the oldest confidence limits for wasp divergence are considered, wasp lineage divergence postdates the earliest host split by more than four million years. This strong temporal asymmetry is inconsistent with strict long-term codivergence and instead supports recent tracking of pre-existing host lineage structure.

Historical effective population size trajectories also differed between the wasp and host lineages (**Fig. 3b and Supplementary Fig. 3b, d**). SMC++ and PSMC inferred changes in wasp effective population size around major climatic intervals, including a decline around one million years ago and a later decline around 100 Kya. Because these *Ne* changes are not central to distinguishing host tracking from codivergence, we treat them as supplementary demographic context rather than as a primary result.

### Host volatile profiles and wasp preferences are lineage-associated

We next tested whether host volatile differentiation and wasp behavioral responses were consistent with olfactory-mediated host recognition. In Y-tube bioassays, wasp choices were strongly lineage-associated: individuals preferentially selected volatiles from plants belonging to their local population or lineage (**Fig. 4a–c**). This behavioral pattern provides direct evidence that wasps discriminate among host lineages using volatile cues and suggests a mechanism by which host fidelity may be maintained at a fine spatial scale.

**Fig. 4.**
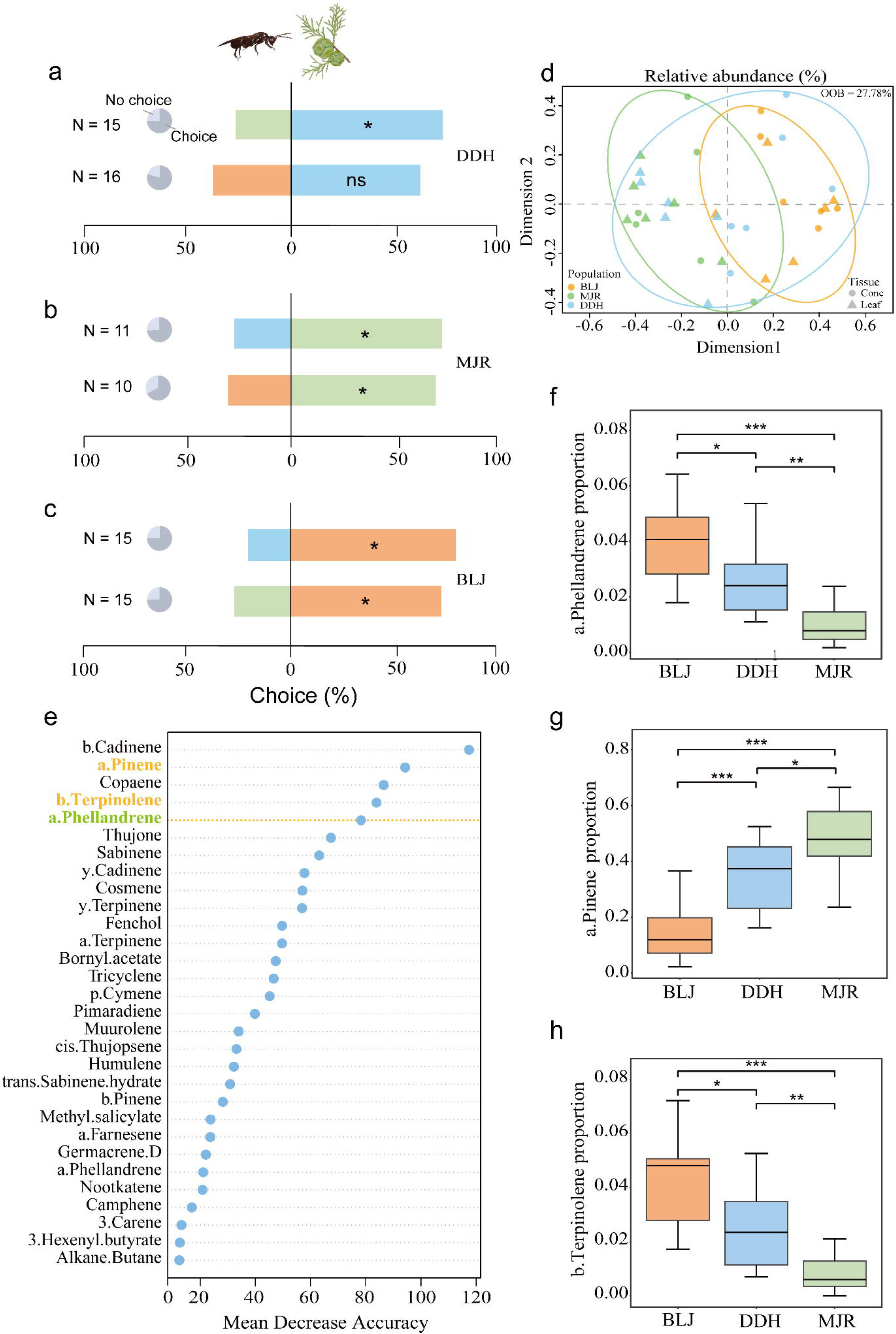
| Effect of host volatile emissions on wasp behavior. **a-c,** Proportion of insects choosing the two odors, where the pie charts illustrate the relative proportion of insects that made or did not make a choice. **a** The plant of DDH population vs. MJR and BLJ. **b,** The plant of MJR population vs. DDH and BLJ. **c,** The plant of BLJ population vs. DDH and MJR. **d, e,** Random Forest (RF) classification analysis of volatile profiles from the three populations of cypress plants (leaf and cone). **d,** Multidimensional scaling plot of RF model classifying volatile bouquets released from different populations. During 100,000 replicated tree constructions samples, with the out-of-bag (OOB) error being 27.78%. **e,** Importance ranking of volatile compounds contributing to separation between different populations plants based on mean decrease in accuracy (MDA) following RF analysis. **f-h,** The difference of volatiles emission proportion (*α*-phellandrene, *α*-pinene and *β*-terpinolene) in different populations of cypress. Asterisks indicate significant differences after T-test or Wilcoxon test (* *p* < 0.05, ** *p* < 0.01, ** *p* < 0.001).

GC-MS analysis of 36 headspace collections from the three host populations detected 61 VOCs, dominated by terpenoids (**Supplementary Table 11**). Multidimensional scaling (MDS) following random forest analysis of proportional VOC data separated the volatile profiles of BLJ and MJR, while DDH encompassed or partially overlapped the variation represented by these groups (**Fig. 4d**). Among 100,000 replicate random forest tree structures, the five most important compounds for discriminating among host populations included three monoterpenes: α-phellandrene, α-pinene, and β-terpinolene, which differed significantly in abundance among populations (**Fig. 4e–h**). Together, the VOC profiling and behavioral assays identify host volatile differentiation as a likely sensory basis for local host recognition, without requiring the stronger claim that VOCs alone caused genome-wide divergence.

### Candidate genomic and phenotypic correlates of host-associated differentiation

We combined gene-environment association analyses with population-specific selective scans to identify genomic regions potentially associated with host-associated and environmental differentiation. However, because the three wasp lineages were sampled from separate river valleys that also differ in altitude, temperature, and precipitation, environmental variation cannot be fully separated from population structure. We therefore treat these loci as candidate signals requiring further validation, rather than as direct evidence for environmentally driven local adaptation.

Using LFMM and RDA, we identified 66 candidate genes associated with environmental variables (**Supplementary Fig. 5a and Supplementary Table 6**). These genes were enriched in 102 GO categories (Weight01 Fisher’s test, *p* < 0.05; **Supplementary Table 7**), including categories related to environmental response, physical development, and metabolic regulation. Candidate genes included *dnc*, associated with heat response; *goa-1*, associated with olfactory adaptation; *ROBO1*, *SLC22A16*, and *sls*, associated with organ development; and *SLC25A25*, associated with food-stimulus response (**Supplementary Fig. 5c and Supplementary Table 8**). Because of the spatial structure of the sampling design, these candidate genes should be viewed as hypotheses for future functional validation.

We also applied XP-CLR and XP-EHH to test for population-specific signatures of selection in the high-altitude DDH population, using BLJ as the reference population. This analysis detected 525 candidate genes in DDH (**Supplementary Fig. 5b and Supplementary Table 6**), enriched in 161 GO categories (Weight01 Fisher’s test, *p* < 0.05; **Supplementary Table 9**), including categories related to morphological regulation, clock regulation, and odor response. Candidate genes included *plexB* and *nAChRb1*, associated with olfactory information processing; *per*, associated with diapause and circadian regulation; and *en2*, *sls*, and *SKP2*, associated with organ development (**Supplementary Fig. 5d and Supplementary Table 10**). These results are consistent with the involvement of sensory, developmental, and phenological pathways in population differentiation, but they should not be interpreted as direct proof of causal adaptation.

Morphological measurements also differed among *M. daduheensis* populations. DDH individuals were larger in body size, ovipositor sheath length, and wing size than individuals from the other two lineages, while MJR individuals had longer bodies and ovipositors than BLJ individuals (**Fig. 5a–c**). Morphological traits covaried with several geographic and climatic variables, including altitude, temperature seasonality, diurnal temperature range, and precipitation-related variables (**Fig. 5d**). However, because these environmental variables are closely aligned with river-valley structure, the observed correlations should be interpreted as associations between morphology and population differentiation, rather than as evidence that individual climatic variables directly caused morphological divergence.

**Fig. 5.**
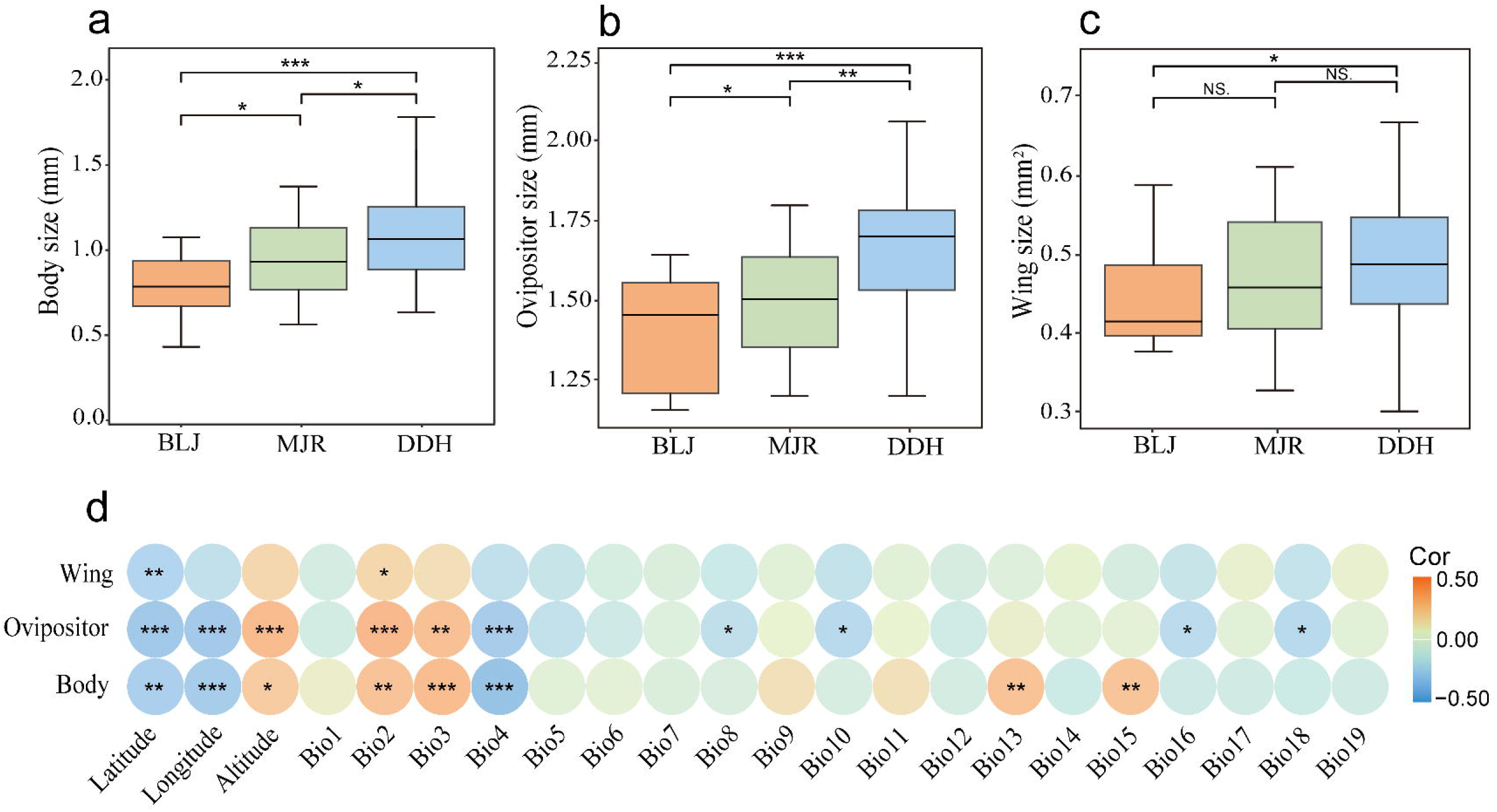
| Differences in morphological characteristics of *M. daduheensis* and the correlation between traits and environment. **a-c,** Differences in body, ovipositor, and wing sizes among the three populations. **d,** Heat map of correlation between three traits and environmental factors.

Together, these results are consistent with recent host-associated differentiation in *M. daduheensis* following the establishment of older host lineage structure. Among the evidence presented here, the VOC profiles and behavioral assays provide the most direct support for olfactory-mediated host recognition.

## Discussion

Genetic differentiation in phytophagous insects is often shaped by host association, environmental heterogeneity, dispersal limitation, and demographic history^22–24^. Our analyses of the *Megastigmus daduheensis* and *Cupressus chengiana* association reveal a phylogeographic pattern that is spatially concordant but temporally asymmetric. Across the Daduhe, Minjiang, and Bailong river catchments, wasp population structure closely mirrors the structure of its host. However, the timing of divergence differs markedly between trophic levels: host lineages diverged during the Pliocene– Pleistocene, whereas point estimates place the major splits among wasp lineages at approximately 7.8–9.1 Kya. Although the confidence intervals for wasp divergence extend into the late Pleistocene, their upper bounds remain several million years younger than the earliest host split. This temporal mismatch is therefore more consistent with rapid host tracking of pre-existing host structure than with long-term synchronous cospeciation^23, 24^.

The young divergence of *M. daduheensis* lineages suggests that population differentiation occurred during a period of late Quaternary environmental reorganization. Similar postglacial demographic responses have been reported in other insects occupying heterogeneous or montane environments^25, 26^. In contrast, *C. chengiana* retained much older lineage structure through Quaternary climatic oscillations, consistent with long-term persistence, allopatric divergence, and secondary gene flow in the Hengduan Mountains and adjacent Qinghai-Tibet Plateau^19-21^. The asymmetry between these histories indicates that the wasp did not simply share the deep evolutionary history of its host. Rather, wasp populations appear to have differentiated after host lineages were already established. Early human activity and habitat modification on the Qinghai-Tibet Plateau may have further altered the distribution and connectivity of suitable habitats during the late Pleistocene and Holocene^27–32^, potentially reinforcing the spatial isolation already imposed by river-valley geography.

Genomic and phenotypic differentiation in *M. daduheensis* is consistent with this recent host-tracking scenario. Selective-scan and gene-environment association analyses identified candidate loci associated with sensory function, development, circadian regulation, and environmental variables. Such patterns are in line with studies showing that adaptation to heterogeneous environments can leave uneven genomic signatures in insect populations^33, 34^. Several candidate genes are linked to developmental or seasonal processes, including circadian regulation, which has been implicated in life-history divergence in other phytophagous insects^35^. These genomic signals correspond with clear phenotypic differences among wasp lineages. DDH individuals were generally larger and had longer ovipositor sheaths and wings than individuals from the other lineages, while MJR and BLJ also differed in body and ovipositor measurements. Comparable associations between morphology, altitude, and thermal environment have been reported in other hymenopteran systems^36–38^. Together, these results indicate that recent population divergence in *M. daduheensis* involved both genomic and morphological differentiation.

Temporal differentiation may further contribute to population separation in this system. Genes associated with circadian or seasonal regulation, together with differences in local climate and host environments among river valleys, suggest that phenological divergence could accompany host-associated differentiation. Allochrony is a recognized route to reproductive isolation in insects, especially where host phenology structures the timing of emergence, mating, or oviposition^39–41^. Although direct phenological measurements are not yet available for *M. daduheensis*, the genomic and environmental patterns observed here make temporal divergence a plausible component of the broader differentiation process.

Olfactory-mediated host recognition provides a mechanistic link between host differentiation and wasp host use. Seed-cone VOC profiles differed among *C. chengiana* lineages, and Y-tube bioassays showed that wasps preferentially responded to volatiles from their local host lineages. Host volatiles are widely used by herbivorous insects during host location and host choice, and variation in volatile blends can influence feeding and oviposition decisions^11, 12^. In *M. daduheensis*, the match between host VOC differentiation and lineage-specific wasp preference indicates that olfactory cues mediate host recognition across river-valley lineages. Candidate genes associated with olfactory adaptation and olfactory circuit development, including *goa-1* and *plexB*, further support the role of sensory differentiation in this process^42, 43^. These chemical, behavioral, and genomic results suggest that host VOCs promote host fidelity and help maintain host-associated differentiation among wasp lineages.

By integrating population genomics, demographic inference, morphology, host volatile profiling, and behavioral assays, this study identifies a distinct evolutionary trajectory for a specialist conifer seed-feeder. The spatial match between *M. daduheensis* and *C. chengiana* does not require synchronous cospeciation. Instead, the wasp lineages appear to have tracked older host lineage structure after host divergence had already occurred. This asynchronous process can generate host–herbivore genetic concordance over a relatively short evolutionary interval. In this system, rapid host tracking, olfactory-mediated host recognition, and host-associated local adaptation together provide a coherent explanation for the emergence of closely matched population structure across trophic levels.

## Methods

### Sample collection and DNA extraction

Female *Megastigmus daduheensis* lay their eggs inside the cones of *C. chengiana*. Each larva feeds on an individual seed, after which it pupates, and the adult chews its way out of the cone (**Fig. 1**)^18, 19^. *Cupressus chengiana* is only found in three isolated arid valleys between 800 and 2,900 m a.s.l. in the upper reaches of the BLJ, DDH and MJR rivers^19^. In 2020–2021 we sampled 11 locations spanning all three river catchments (**Fig. 1**), collecting seed cones from 15 trees from each location (**Fig. 1 and Supplementary Table 1**). Adult wasps were reared from the cones in mesh bags, and on emergence they were immediately frozen in liquid nitrogen and preserved at -80°C until DNA extraction.

We used 216 individual wasps for whole-genome resequencing, including 211 *M. daduheensis* individuals from the three focal river catchments and five individuals of the closely related species *M. duclouxiana* as outgroups. The *M. daduheensis* samples comprised 97 individuals from DDH, 60 from MJR, and 54 from BLJ. Genomic DNA was extracted using the DNeasy Blood & Tissue Kit (Qiagen), following the manufacturer’s protocol. DNA concentration and integrity were assessed using the Qubit® DNA Assay Kit on a Qubit® 3.0 Fluorometer (Invitrogen, USA) and agarose gel electrophoresis.

### Sequencing, Read Mapping, and Quality Control

Genomic library preparation was performed by Novogene (Beijing, China) and sequenced with 18× coverage on the Illumina platform with 150 bp paired-end reads. We used Fastp v0.20.0 ^43^ to perform basic statistics on the quality of the raw reads and to filter out low quality reads pairs, including reads with >50% of low quality bases (Q≤5) in either of the paired reads, and reads containing >10% Ns. Subsequently, clean reads were mapped to the reference genome of *M. duclouxiana*^44^ using the BWA-MEM v0.7.17^45^. The resulting bam files were sorted using samtools v1.10^46^, and duplicated reads were removed by sambamba v0.7.1^47^. The sequencing and mapping statistics are summarized in **Supplementary Table 2**.

SNP calling was performed using the GATK v4.2.0.0^48^ HaplotypeCaller. SelectVariants was first used to extract SNP sites. VariantFiltration was then used to marker and filter the SNPs. The filtering settings were as follows: DP < 5 | | QD < 2.0 | | MQ < 40.0 | | FS > 60.0 | | SOR > 3.0 | | MQRankSum < −12.5 | | ReadPosRankSum <−8.0. Finally, VCFtools v0.1.17^49^ was used to filter the SNPs which had a missing > 20 (--max-missing 0.8) and minor allele frequency (MAF) < 5% (--max-missing 0.05).

The 53 transcriptome datasets for host *C. chengiana* were obtained from a previous study by our group^19^, which included 11 populations in the same three river catchments sampled for the wasps. Sampling comprised 29 samples from DDH, 10 from MJR, 14 from BLJ; four samples of *Cupressus funebris* were included as outgroups.

### Population Structure

We used a combination of population structure analysis, principal component analysis (PCA), and phylogenetic analysis of the SNP data sets to evaluate the genetic structuring of *M. daduheensis* and *C. chengiana* in the three river catchments. Firstly, VCFtools v0.1.17 and Plink v1.9^50^ were used to convert input data and remove linkage disequilibrium (LD) sites with the parameter set as “--indep-pairwise 50 5 0.2.” Then, ADMIXTURE v1.23^51^ was used to estimate the population structure with *K* (the number of genotype clusters) set to 2–7. The minimization of cross-validation error among all runs was used to determine the most likely value of *K*. We also explored the genetic structure of each species using PCA implemented in Plink v1.9^50^. We inferred maximum-likelihood (ML) phylogenetic trees for each species using IQ-TREE v2.1.3^52^ with *M. duclouxiana* and *C. funebris* as outgroups for the wasp and conifer phylogenies, respectively. Nodal support values were obtained for 1000 bootstrap calculations.

Finally, we also used VCFtools v0.1.17 to estimate the value of Tajima’s *D*, nucleotide diversity (π) and population genetic differentiation (*F_ST_*). LD analysis was performed using the PopLDdecay software^53^. The SNP spacing on the genome was fitted to the linkage disequilibrium coefficient (*r*^2^), and data with *r*^2^ greater than 0.1 were used as the decay value of the LD.

### Demographic history

We estimated divergence times using *fastsimcoal2* (FSC2)^54^ based on fourfold degenerate synonymous sites (4DTv) to reduce the influence of natural selection. Absolute time calibrations require assumptions about mutation rates (μ) and generation times (g). For *M. daduheensis*, we applied a mutation rate of 3.6 × 10⁻⁹ per site per generation, derived from direct estimates in bumblebees^55^ (the closest available phylogenetically relevant insect with published mutation rate estimates), and a generation time of one year based on field observations of univoltine life history. For *C. chengiana*, we used a mutation rate of 9.7 × 10⁻⁹ per site per generation and a generation time of 50 years^19^. We emphasize that absolute time estimates carry substantial uncertainty (reflected in 95% confidence intervals; see Results), and that the most robust conclusion is the relative difference between trophic levels, which spans orders of magnitude, rather than the precise calendar ages. Specifically, even if wasp divergence times were at the upper bounds of their 95% CIs (∼167 Kya), they would still postdate host divergence (∼4.56 Mya) by more than 4 million years. The same qualitative pattern (Pliocene host divergence vs. Holocene wasp differentiation) holds across a wide range of plausible mutation rates and generation times (sensitivity analysis in **Supplementary Text S1**).

To reduce the potential influence of natural selection, only SNPs located at fourfold degenerate synonymous (4DTv) sites were retained for demographic inference. Both variable and invariant callable sites were retained when constructing the site-frequency spectra, and no minor-allele-frequency filter was applied for demographic analyses. SNPs without minor allele frequency filtering were further filtered to remove sites with any missing data across all sampled individuals. A folded two-dimensional joint site frequency spectrum (2D-SFS) was constructed using a custom Perl script. Based on the genetic structure of both the wasp populations and their host plants, we formulated eight alternative evolutionary models, which are illustrated in **Supplementary Fig. 4**. These models fall into three major categories: (a) divergence of the MJR and BLJ lineages from a recent common ancestor [((MJR, BLJ), DDH), models1–3]; (b) divergence of the MJR and DDH lineages from a recent common ancestor [((MJR, DDH), BLJ), models 4–6]; and (c) radiative evolution among all three lineages [(MJR, DDH, BLJ), models 7–8]. In each scenario, different forms of gene flow were incorporated, resulting in a total of eight models. Details of the model parameter settings, mutation rates, generation time assumptions, and bootstrap procedures are provided in the Supplementary Information (**Supplementary Text and Supplementary Fig. 4**).

We selected one individual with high sequencing depth from each group to estimate the demographic history of *M. daduheensis* using Pairwise Sequentially Markovian Coalescence^56^ (PSMC). The parameters were set as -N25 -t15 -r5 -p “4+25*2+4+6”, with a mutation rate of 3.6e^-9^ based on bumble bees^55^, and the generation time of *M. daduheensis* was determined as one year based on our field observations and previous publications. SMC++^57^, a recently published approach with higher resolution for the recent past when compared to PSMC, was also used to predict the demographic history (population sizes and divergence times) of *M. daduheensis* based on multiple unphased female individuals. Three groups (or lineages) previously defined based on population structure were used for the analysis. The short generation time of *M. daduheensis* allows for reliable and precise estimation of effective population sizes (*Ne*) and divergence times in the recent past using the method of SMC++.

The only new demographic analysis for the host plant *C. chengiana* was a re-estimation of *Ne* with SMC++, with a mutation rate of 9.7e^-9^ and a generation time of 50 years^19^. The aim was to compare this estimate with previous analyses from our group using Stairway Plot^19^.

### Dual-choice olfactory bioassays

The plants and wasps used in this part of the study were collected between 17–23. Oct. 2023. Detailed methods are provided in the supplemental material (**Supplementary Fig. 6**). To assess whether *M. daduheensis* adults can use plant volatiles as host location cues, we conducted Y-tube olfactometer bioassays to test their olfactory responses to VOCs emitted by plants from three river catchments (DDH, MJR, and BLJ).

For each population, eight to fifteen trees were sampled, and three to five cone-bearing branches were collected per tree. For each trial, fresh leaves or cones originating from a single population were pooled and placed into one odor source bottle, such that each arm of the Y-tube contained plant material from a different population. The following population pairs were compared: (a) DDH vs. MJR, (b) DDH vs. BLJ, (c) MJR vs. DDH, (d) MJR vs. BLJ, (e) BLJ vs. DDH, and (f) BLJ vs. MJR. Odor sources were alternated between arms every five trials to control for positional bias.

Dual-choice bioassays were conducted using glass Y-shaped olfactometers (15.0 cm base arm length, 15.0 cm side arm length, 3.0 cm internal diameter). Newly emerged adult wasps, starved for 12–24 h, were individually tested, with each insect used in only one trial (n = 82). A wasp was recorded as making a choice when it walked at least two-thirds of the length of one side arm and remained there for at least 30 s; individuals that did not make a choice within 5 min were scored as “no choice”.

### VOCs collection and analysis

We selected six plants from each of three populations (n=18) for volatile collection, and collected leaves and cones separately for each individual for a total of 36 samples. VOCs were collected using a push–pull headspace sampling system. The incoming air was maintained as it was during the olfactory bioassay, with a flow rate of about 300 mL min^−1^. The headspace was pulled out with a vacuum pump at 300 mL min^−1^ through a stainless-steel adsorbent cartridge filled with Tenax TA (150 mg of each, mesh 60; Markes International, Llantrisant, RCT, UK). Volatiles were collected for 15 min.

Volatile samples were analyzed by gas chromatography–mass spectrometry (GC–MS; Agilent GC 7890A, MSD 5975C, New York, USA). Trapped compounds were thermally desorbed (TD100, Markes International, Llantrisant, RCT, UK) at 250 ◦C for 10 min, cryofocused at 10 ◦C and injected onto an HP-5 capillary column (50 m × 0.2 mm; film thickness 0.5 µm) with helium as a carrier gas at a flow of 1.2 mL min^−1^. The column temperature was held at 40 ◦C for 1 min, then ramped at 5 ◦C min^−1^ to 210 ◦C, and ramped again at 20 ◦C min^−1^ to 250 ◦C. Chromatograms were analyzed using the software PARADISe v3.9 ^58^. Individual VOCs were tentatively identified by comparing mass spectra to those in the NIST 2014 mass spectral library, verified by chromatography with authentic standards where available and quantified based on external standards. For those compounds that had no authentic standards available, quantification was estimated based on α-pinene.

### Statistical analysis

All statistical analyses were performed in R v4.1.3^59^. We performed statistical analysis of variation in body size, fore wing area, and ovipositor sheath length across wasp populations sampled from the three river catchments (i.e. DDH, MJR and BLJ). Prior to analysis all datasets were tested for normality using the Shapiro-Wilk test in R. We used a paired-sample *T*-test or a paired-sample Wilcoxon test in the ggsignif^60^ package to test for significant differences in traits between populations. We then investigated correlations between three traits and environmental factors using the “cor.test” function, which calculates a Pearson’s product moment correlation coefficient.

During the dual-choice bioassays, differences between the two odor fields were assessed using *T*-test or Wilcoxon test, again after assessing whether the data were normally distributed. In addition, the “ggsignif” package was also used to test the difference of volatile emissions between plants of the three populations.

To characterize differences in volatile blends among populations from the three river cathments, we used a RF between-group classification algorithm as described previously^61^. RF analysis is similar to a principal component analysis, but is more suitable for nonparametric datasets that comprise many more variables than samples^61^. We performed RF analysis on the emission rates of individual compounds of the entire blend by using the package randomForest in R. For each analysis, 100,000 bootstrap samples were drawn with 10 variables (i.e., volatiles) randomly selected at each node. The number of variables selected is based on the square root of all variables. Differences among populations were then visualized using MDS plots. The importance of each compound for the distinction is expressed as the mean decrease in accuracy (MDA) and the odds of the compound being improperly classified is expressed as the out-of-bag (OOB) error rate.

### Compilation of climate and altitude data

Climate layers comprising 19 bioclimatic variables (**Supplementary Table 12**) were retrieved for current climates (1970–2000) at 30 seconds resolution from the WorldClim database^62^(https://www.worldclim.org/), along with one altitude layer was downloaded from the SRTM elevation database (https://www2.jpl.nasa.gov/srtm/). The Spatial Analysis tool in ARCGIS v.10.7 (Environmental Systems Research Institute, Redlands, CA, http://www.esri.com) was used to extract the climate and altitude data for each layer respectively corresponding to the geographical coordinates each of the 11 locations. To avoid multicollinearity, we calculated pairwise Pearson’s correlation coefficients^63^ (*r*) for current climate and altitude data in R v4.1.3^59^, then excluded any factor that had a correlation coefficient greater than |0.8| with two or more other factors. We randomly eliminated one out of each pair of correlated factors, ultimately retaining a subset of six uncorrelated BIOCLIM variables for subsequent GEA (Gene-environment association) analysis: Bio3 (Isothermality), Bio7 (Temperature Annual Range), Bio10 (Mean Temperature of the Warmest Quarter), Bio11 (Mean Temperature of the Coldest Quarter), Bio12 (Annual Precipitation), and Bio15 (Precipitation Seasonality).

### Local adaptation and selective sweeps in *M. daduheensis*

Two methods were used to measure the associations between all SNPs and climatic gradients: latent factor mixed model^64^ (LFMM) and redundancy analysis^65^ (RDA), representing univariate and multivariate GEA approaches, respectively. Further details on these two methods are provided in the supplementary materials.

To explore the adaptation of wasps to high altitude, we used XP-CLR and XP-EHH to compare the selection signal in the higher altitude DDH population of *M. daduheensis* with the lower altitude BLJ population. XP-CLR is a method for the determination of the selection signal based on differentiation of allele frequencies across populations, which can avoid ascertainment biases and successfully detect older signals and selections on standing variation^66^. The putative selection targets were extracted as the highest 5% value for XP-CLR, with a sliding window of 10 kb. Furthermore, we also computed the XP-EHH values using haplotype information in the XP-EHH program to determine the positive selection in DDH population^67^. To detect positive selection, we split the genome into non-overlapping segments of 10 kb and computed the maximum XP-EHH score in each segment. Regions with the highest and lowest 1% of XP-EHH were considered as candidate regions in DDH.

Finally, we performed a Gene Ontology (GO) enrichment analysis using the Bioconductor package TOPGO v2.24.0^68^ to identify the biological processes associated with the genes identified as candidates for local adaptation outliers.

### Morphological measurements

We measured body size, fore wing area, and ovipositor sheath length (as a proxy for ovipositor length) using ImageJ^69^ on photos of 122 female samples of *M. daduheensis*. The wasps were removed from the vial, gently pressed prone to the slide with the pronotum parallel to the digital microscope lens using flat forceps and the dorsal view was taken. Then, we flattened out the fore wing and ovipositor sheath to capture the entire lateral view. The body size of the wasps was estimated using thoracic inter-tegular distance^70^. During image acquisition, all specimens were photographed under the same standard and scale.

## Acknowledgements

We thank Dr. Jianquan Liu for his constructive comments. This project was funded by the National Natural Science Foundation of China (grants U25A20644), the Fundamental Research Funds for the Central Universities of Sichuan University (grants SCU2025D003, SCU2024D003).

## Arthur Contributions

XXC designed research; XXC performed research; XXC, JGK, YW, JLL, HLL, TL and SLY analyzed data; XXC, JGK, WQH, WTW collected data; YMZ, GNS, and KSM reviewed & edited the paper; XXC, YMZ, GNS, and KSM wrote the paper.

## Data Availability

All data needed to evaluate the conclusions in the paper are present in the paper and/or the Supplementary Materials. All sequencing data (whole-genome resequencing data for 211 individuals) in this study have been deposited in the National Center of Biotechnology Information (https://www.ncbi.nlm.nih.gov/) under accession number SUB14560533.

---

**Supplementary Fig. 1** | **Results of cross-validation (CV).**

**Supplementary Fig. 2** | **Genetic parameter of *M. daduheensis* and *C. chengiana*. a, b,** Paired genetic distances (*F_st_*) between individuals from three lineages. **c, d,** Nucleotide diversity (*π*) of three lineages. **e,** The results of LD (expressed as *r*^2^) in the wasps showed that DDH population had the fastest decay rates, indicating that it had the lowest genetic diversity.

**Supplementary Fig. 3** | **Demographic history of *M. daduheensis* and *C. chengiana*. a,** Phylogenetic relationships and divergence times are shown for *M. daduheensis* with other *Megastigmus* species. **b,** Changes in effective population size (*Ne*) through time inferred by the Pairwise Sequentially Markovian Coalescent (PSMC) model. **c,** Divergence times of wasp populations inferred by SMC++. The differentiation time of DDH and BLJ was 12.3 Kya, DDH and MJR was 10.5 Kya, and BLJ and MJR was 4.2 Kya. **d,** Historical effective population size of hosts based on re-sequencing data predicted by SMC++.

**Supplementary Fig. 4** | Schematic diagram of 8 tested demographic models using *fastsimcoal27*.

**Supplementary Fig. 5** | **Genomic variants associated with the three populations of *M. daduheensis*. a,** A brief framework for GEA analysis. The Venn diagram shows the number of overlapping genes identified by RDA and LFMM, and the number of genes identified by temperature and precipitation. **b,** A brief framework for selective sweeps analysis. The Venn diagram shows the number of overlapping genes identified by XP-CLR and XP-EHH. **c, d,** The scatter plot represents the GO categorization of identified genes in significant pathways in GEAs and selective sweeps analysis, respectively (*p* < 0.05). The color of the circle shows the −log (*p*) value. The size of the circle shows the number of potential environment-associated genes in the enriched pathway. **e,** The scatter plot represents the GO categorization of identified genes in significant pathways in temperature (*p* < 0.05). The color of the circle shows the -log (*p*) value. The size of the circle shows the number of potential environment-associated genes in the enriched pathway. **f,** The scatter plot represents the GO categorization of identified genes in significant pathways in precipitation (*p* < 0.05). The color of the circle shows the -log (*p*) value. The size of the circle shows the number of potential environment-associated genes in the enriched pathway.

**Supplementary Fig. 6** | **The entire experimental setup of olfactory bioassay.**

